# MHC genotyping from rhesus macaque exome sequences

**DOI:** 10.1101/625244

**Authors:** John R. Caskey, Roger W. Wiseman, Julie A. Karl, David A. Baker, Taylor Lee, Muthuswamy Raveendran, R. Alan Harris, Jianhong Hu, Donna M. Muzny, Jeffrey Rogers, David H. O’Connor

**Author notes:** Corresponding author: Dr. David H. O’Connor.

## Abstract

Indian rhesus macaque major histocompatibility complex (MHC) variation can influence the outcomes of transplantation and infectious disease studies. Frequently, rhesus macaques are MHC genotyped to identify variants that could account for unexpected results. Since the MHC is only one region in the genome where variation could impact experimental outcomes, strategies for simultaneously profiling variation in the macaque MHC and the remainder of the protein coding genome would be useful. Here we introduce <u>m</u>acaque <u>e</u>xome <u>s</u>equence (MES) genotyping, in which MHC class I and class II genotypes are determined with high confidence using target-enrichment probes that are enriched for MHC sequences. For a cohort of 27 Indian rhesus macaques, we describe two methods for obtaining MHC genotypes from MES data and demonstrate that the MHC class I and class II genotyping results obtained with these methods are 98.1% and 98.7% concordant, respectively, with expected MHC genotypes. In contrast, conventional MHC genotyping results obtained by deep sequencing of short multiplex PCR amplicons were only 92.6% concordant with expectations for this cohort.

## Introduction

The major histocompatibility complex (MHC) is an intensively studied set of genes in macaques (Shiina et al. 2017; Wiseman et al. 2013). The genomic MHC region contains clusters of genes that encode the MHC class I complex and the MHC class II complex. Cells use MHC class I molecules to present intracellular peptides to immune cells like the CD8+ T cell or natural killer cells (Garcia and Adams 2005). MHC class I molecules accommodate intracellular peptides of varying specificity by having diverse amino acid sequences in the *α*1 and *α*2 subunits, which form the peptide-binding cleft (Silver and Watkins 2017; Loffredo et al. 2009). These *α* 1 and *α* 2 subunits correspond to exons 2 and 3, respectively, of an MHC class I gene (Malissen et al. 1982). Most of the polymorphisms that distinguish MHC class I alleles are concentrated in exons 2 and 3 (Williams 2001). Thousands of individual MHC allelic variants have been identified in the three most widely used macaque species for biomedical research: rhesus (*Macaca mulatta*), cynomolgus (*Macaca fascicularis*), and pig-tailed macaques (*Macaca nemestrina*) (Semler et al. 2018; Karl et al. 2017; Maccari et al. 2017).

In humans, the MHC is also termed the human leukocyte antigen complex (HLA). HLA class I has a single copy of the *HLA-A, HLA-B*, and *HLA-C* genes on each chromosome (Daza-Vamenta et al. 2004; Shiina et al. 2017). In contrast, macaques have a variable number of genes on each chromosome that encode MHC class I MHC-A and MHC-B proteins, and macaques lack an *HLA-C* orthologue (Daza-Vamenta et al. 2004; Shiina et al. 2017; Wiseman et al. 2013). Initial approaches to genotype macaque MHC class I relied on using sequence-specific PCR oligonucleotides to test for the presence or absence of individual alleles (Kaizu et al. 2007). More recently, deep sequencing genomic DNA or complementary DNA PCR amplicons spanning a highly variable region of exon 2 has become commonplace (Wiseman et al. 2009; Karl et al. 2013). Amplicon sequences can be used to genotype groups of closely related MHC class I alleles, which are denoted as lineages. For example, an amplicon deep sequence that corresponds to the rhesus macaque *Mamu-A1*001* lineage demonstrates that an animal possesses *Mamu-A1*001:01*, *Mamu-A1*001:02*, or another closely related variant that has not yet been identified. This lineage-level reporting of MHC class I genotypes can be sufficient for designing experiments where certain MHC class I genotypes need to be matched between animals, balanced among experimental groups, or excluded entirely from a study (Loffredo et al. 2007; Loffredo et al. 2009; Muhl et al. 2002; Karl et al. 2013; Wiseman et al. 2009; Nomura et al. 2012; Mothe et al. 2003).

Amplicon deep sequencing can also be used for MHC class II genotyping. The human HLA class II genes *DQA1, DQB1, DPA1*, and *DPB1* have direct orthologues in macaques (Otting et al. 2017). Both macaques and humans have a variable number of MHC class II DRB genes on a single chromosome, while the DRA gene is oligomorphic (Daza-Vamenta et al. 2004; Shiina et al. 2017), and typically is not used for genotyping purposes. MHC class II molecules are members of the immunoglobulin superfamily, but they differ from MHC class I in several ways. The MHC class II complex is comprised of an α and β subunit heterodimer, and separate genes encode each MHC class II α and β subunit (Brown et al. 1993). Highly polymorphic regions that are diagnostic for MHC class II allele lineages can be PCR amplified in a manner that is similar to the MHC class I genotyping. The most extensive polymorphism among alleles of the MHC class II genes is found in exon 2 (Williams 2001). Each of the *DRB, DQA1, DQB1, DPA1* and *DPB1* polymorphic MHC class II genes are sufficiently divergent that separate PCR amplicons are required for deep sequencing (Karl et al. 2014).

Comprehensive MHC class I and class II genotyping of macaques by amplicon deep sequencing requires preparation of six separate PCR amplicons for MHC class I and class II DRB, DQA1, DQB1, DPA1, and DPB1 (Karl et al. 2014; Karl et al. 2017). Despite this complexity, a major advantage to amplicon deep sequencing has been its cost effectiveness. The output from a single MiSeq sequencing run can be used to determine the MHC class I and MHC class II genotypes for up to 192 macaque samples. In recent years, improved sequencing hardware and software have prompted new approaches to MHC genotyping. Instead of a utilizing PCR amplicons of variable gene regions to genotype samples, researchers can use human <u>w</u>hole <u>e</u>xome <u>s</u>equencing (WES) and <u>w</u>hole <u>g</u>enome <u>s</u>equencing (WGS) datasets to determine HLA genotypes (Xie et al. 2017; Kishikawa et al. 2019; Yang et al 2014; Posey et al. 2016). Likewise, target capture approaches have been described for HLA genotyping with next-generation sequencing (Cao et al. 2013; Wittig et al. 2015). In contrast, MHC genotyping of macaques from WGS (Xue et al. 2016; Bimber et al. 2017; de Manuel et al. 2018) or whole exome sequence (Vallender 2011; Cornish et al. 2016) datasets have not been reported to date. This reflects the challenges presented by mapping short sequence reads against complex, duplicated gene families like the MHC class I genes of macaques where genomic reference sequences and reference databases of known alleles are incomplete (Zimin et al. 2014; Maccari et al. 2017). In an era where the per-base cost of sequencing is dropping rapidly, we explored the feasibility of obtaining whole exome sequencing data while maintaining parity of results with the traditional MHC PCR amplicon approach.

Here we introduce MHC genotyping via macaque exome sequencing (MES), which is an exome sequencing-based workflow for comprehensive MHC class I and class II genotyping in macaques. This workflow uses a commercially available human exome target-capture enrichment kit in conjunction with specialized spike-in target-capture probes to specifically cope with the high copy number of macaque MHC genes. We show that accuracy of Indian rhesus macaque MHC class I and class II results from this workflow are comparable to conventional MiSeq genotyping when using exon 2 reference sequences.

## Methods

### Animals

Twenty-seven whole blood samples were collected from Indian rhesus macaques (*Macaca mulatta*). Five of these samples came from a breeding group of animals living at the Wisconsin National Primate Research Center (WNPRC). The remaining 22 samples were provided by Dr. Michele Di Mascio from the National Institutes of Health’s National Institutes of Allergy and Infectious Diseases. Blood sampling was performed under anesthesia and in accordance with the regulations and guidelines outlined in the Animal Welfare Act, the Guide for the Care and Use of Laboratory Animals, and the Weatherall report (Animal Welfare Act 1966; Weatherall 2006).

### Data

Exome sequence datasets have been deposited in the sequence read archive (SRA) under BioProjects PRJNA529708 and PRJNA527214. Fasta reference sequences used for MHC genotyping, and sequence analysis scripts are available from https://go.wisc.edu/jb0926.

### MHC class I and class II genotyping by amplicon deep sequencing

Genomic DNA was isolated from 250*μ*L of whole blood using a Maxwell^®^ 48 LEV Blood DNA Kit (Promega Corporation, Fitchburg, WI). Following isolation, DNA concentrations were determined with a Nanodrop 2000 and samples were normalized to 60 ng/ul. MHC class I and class II PCR amplicons were generated using exon 2-specific primers with adapters (CS1 and CS2) necessary for 4-primer amplicon tagging with the Fluidigm Access Array™ System (Fluidigm, San Francisco, CA, USA) by previously described methods (Karl et al. 2017; Karl et al. 2014). Pooled PCR products were purified using the AMPure XP beads (Agencourt Bioscience Corporation, Beverly, MA, USA) and quantified using the Quant-iT dsDNA HS Assay kit with a Qubit fluorometer (Invitrogen, Carlsbad, CA, USA), following the manufacturer’s protocols. The MHC exon 2 genotyping amplicon pools were sequenced on an Illumina MiSeq instrument (San Diego, CA, USA) as previously described (Karl et al. 2017).

Analysis of the MiSeq exon 2 genotyping amplicon sequences was performed using a custom Python workflow. The workflow contained a step to remove oligonucleotide primers and sequencing adapters with bbduk, merge a step to reads bbmerge, a step to identify unique sequences/remove chimeras with USEARCH, and finally mapping unique reads against a deduplicated reference database of rhesus macaque partial MHC class I and class II exon 2 sequences with bbmap (Bushnell et al. 2017; Edgar 2010). The database was derived from sequences in the IPD-MHC NHP database downloaded from the European Bioinformatics Institute website (Maccari et al. 2017). For this publication, we define “IPD exon 2” as this database of reference sequences. SAM output files from bbmap were parsed with the Python package pandas to enumerate the reads from each animal that were identical to IPD exon 2 reference sequences. *Mamu-A, -B, -DRB, -DQA1, -DQB1, -DPA1* and -*DPB1* lineage-level haplotypes were inferred for each of the samples with a semi-automated custom workflow that identifies diagnostic alleles associated with previously defined rhesus macaque haplotypes (Karl et al. 2013; Otting et al. 2017).

### MHC class I and class II genotyping by exome sequencing

Genomic DNA was isolated as described above and shipped to the Human Genome Sequencing Center at the Baylor College of Medicine. MHC and exon-containing genomic DNA fragments were selectively enriched using a custom target-capture probeset. Genome-wide exons were captured using SeqCap EZ HGSC VCRome2.1, an optimized human clinical exome probeset (Clark et al. 2013, Yang et al. 2013). SeqCap EZ HGSC VCRome2.1 contains probes designed to enrich 23,585 human genes and 189,028 non-overlapping exons. A low coverage audit was performed to identify rhesus macaque exons inferred from the reference genome rheMac2 that were not sufficiently enriched (<20x coverage) with these human probes, and an additional 22,884 rhesus macaque exons were incorporated into the genotyping probe design (Prall et al. 2017). Finally, and most importantly for MHC analyses, we modified the SeqCap EZ Design: Human MHC Design to selectively enrich MHC class I and class II sequences. This previous design was prepared by the Beijing Genome Institute in collaboration with Roche/Nimblegen and it targeted the complete 4.97Mb HLA region with non-redundant probes designed against 8 fully sequenced HLA haplotypes (Horton et al. 2008; Cao et al. 2013). For our macaque studies, we prepared a minimal MHC target capture design using a subset of these probes that are based on all functional HLA class I (*HLA-A, -B, -C*, and -*E*) and class II (*HLA-DRA, -DRB1, -DRB3, -DRB4, -DRB5, -DQA1, -DQB1, -DPA1* and -*DPB1*) genes. Probes were included to capture complete gene sequences (exons + introns + 3’ UTR) as well as ~1 kb of 5’ upstream flanking sequence. The BED file of rhesus rheMac2 target coordinates lifted over to rheMac8 was used to prepare this combined minimal MHC and supplemental rhesus spike-in probe design. Because derivation of MHC results from genotyping is paramount, we used a ratio of 2.5x spike-in probes to 1x VCRom2.1 probes. The supplemental probes for MHC and rhesus macaque were a single reagent, and the MHC-specific probes only constituted 609 kb of the 37.9 Mb probes (Prall et al. 2017).

An Illumina paired-end pre-capture library was constructed with 750 nanograms of DNA, as described by the Baylor College of Medicine Human Genome Sequencing Center. Pre-capture libraries were pooled into 10-plex library pools for target capture according to the manufacturer’s protocol. Samples were pooled in 10-plex sequence capture library pools for 151 bp paired-end sequencing in a single lane of an S4 flow cell on an Illumina Novaseq 6000 at the Baylor College of Medicine Human Genome Sequencing Center.

### Enumeration of MHC reads in exome data

The effective enrichment of MHC reads in the exome datasets was calculated by mapping the reads of each sample’s exome dataset against an individual genomic reference file for each individual locus: *HLA-A* exons 2-3 (NCBI Gene ID: 3105) and *HLA-E* exons 2-3 (Gene ID: 3133); *HLA-DPA1* exons 2-4 (Gene ID: 3113); *HLA-DPB1* exons 2-4 (Gene ID: 3115); *HLA-DQA1* exons 2-4 (Gene ID: 3117); *HLA-DQA2* exons 2-4 (Gene ID: 3118); *HLA-DQB1* exons 2-4 (Gene ID: 3119); *HLA-DQB2* exons 2-4 (Gene ID: 3120); *HLA-DRB1* exons 2-4 (Gene ID: 3123), *HLA-DRB3* exons 2-4 (Gene ID: 3125), *HLA-DRB4* exons 2-4 (Gene ID: 3126) and *HLA-DRB5* exons 2-4 (Gene ID: 3127). This mapping was done by using bbmap with default parameters, which corresponds to a minimum alignment identity of approximately 76% (Bushnell et al. 2017). Empirically, these mapping parameters are sufficient to map macaque MHC reads to their human orthologues. Mapped reads were written to a new fastq file using bbmap’s outm= parameter. To quantify the total number of reads in a sample and the number of reads extracted with our reference file, we created a custom Python script, which is available to download.

### MHC genotyping from exome data

Two complementary data analysis strategies were employed to analyze the exome sequence data, and to verify reproducibility and confidence in the MHC genotyping results. For accuracy and quantification purposes, the expected MHC genotypes for each animal were established based on concordance among at least two out of the three described strategies and biological plausibility, e.g., no more than two alleles per *Mamu-DQA1, -DQB1, - DPA1*, or*-DPB1* locus.

#### Strategy 1: MHC genotyping using Diagnostic Sub-Region (DSR)

The Diagnostic Sub-Region (DSR) was an intra-allelic region that encompassed polymorphisms, and these polymorphisms were distinguishable from alleles with similar sequences. Therefore, this method ensured the DSR was captured in at least one read for each called allele. MHC class I and class II reads initially were extracted from each animal by mapping the FASTQ reads to HLA class I and class II reference sequences containing exons 2-3 and exons 2-4, respectively, plus the intervening intron(s) as described above in ‘Enumeration of MHC reads in exome data’. Reads were mapped to these reference sequences using bbmap with default parameters and the parameter (qtrim=lr) (Bushnell et al. 2017).

Following extraction of MHC reads from the total exome sequences, the MHC reads were prepared for assembly using a modified version of a data pre-processing pipeline, which included tools from the BBTools package (Bushnell et al. 2017). Briefly, optical duplicates and reads from low-quality regions of the sequencing run were removed. Next, Illumina sequencing adapters were trimmed from the ends of sequencing reads. Any residual spike-in or PhiX sequences that inadvertently survived mapping to HLA class I and class II were then removed. Three rounds of error-correction and read merging were performed to create high-confidence merged reads that were well-supported by common kmers found in the extracted MHC reads. The error-corrected reads were not merged with a minimum overlap, but instead were separately mapped against the IPD exon 2 sequences using bbmapskimmer. The default settings for the software tool bbmapskimmer were used with the following modified parameters (semiperfectmode=t ambiguous=all ssa=t maxsites=50000 maxsites2=50000 expectedsites=50000) (Bushnell et al. 2017). The semiperfectmode setting accepted reads with perfect matches, as well as reads that extended off the end of contigs for no more than half of the length of the mapped read segment. The ambiguous setting and the ‘expectedsites’ setting reported the first 50,000 matched read segments that met the ‘semiperfectmode’ filtering, which was set exceedingly high in order to exhaustively map our IPD exon 2 reference file of ~1600 alleles. Using semiperfect mode, these were the segment sequences with the longest matching length. This output file included all mapped reads to all IPD exon 2 reference allele sequences.

We then used samtools mpileup with the settings (-A -a --ff UNMAP -x -B -q 0 -Q 0) on the bbmapskimmer output to calculate a depth of coverage at each position for each IPD exon 2 sequence. Next, we removed any aligned reads to the IPD exon 2 database sequences that contained less than a minimum depth of coverage of two across the entire reference sequence. For ambiguously-mapped reads, the alignments with the longest-matching region were selected for further analysis; ties among alignments were counted multiple times. To reduce the number of false positives, we used Python pandas to only report database sequence matches that had at least one unambiguously mapped read. Based on these mapping parameters, unambiguously-mapped reads must span the DSR. The MHC genotypes from this method were reported as the minimum depth of coverage for each IPD exon 2 database sequence per animal.

#### Strategy 2: De novo reconstruction of MHC sequences from exome reads

Most *de novo* sequence assemblers have been optimized for resolving long contigs, and are tolerant of small sequence mismatches that can otherwise fragment assemblies. In the case of MHC alleles, however, such closely related sequences were often biologically distinct. Because of the gene duplications in the macaque MHC, there were many valid MHC contigs that could be assembled from a single sample within the exome data. This is conceptually similar to the computational challenge of assembling viral haplotypes, where rapidly evolving viruses such as human immunodeficiency virus accumulate variants that frequently co-segregate as minor populations within an infected person (Baaijens et al. 2017). Therefore, we utilized the overlap assembly algorithm SAVAGE, originally designed to reconstruct viral haplotypes, to reconstruct MHC allele sequences from exome reads (Baaijens et al. 2017). Similar to Strategy 1 above, HLA-mapped reads were pre-processed using the BBTools package to remove low quality reads and optical duplicates. Next, adapters were trimmed, residual spike-in and PhiX sequences were removed, three steps of error-correction took place, and reads were merged. These merged reads, as well as high-confidence unmerged paired-end reads that could not be grouped into an overlapping merged read, were used for SAVAGE assembly. The following workflow was implemented in a reproducible snakemake workflow that fully documents parameter selection (Koster and Rahman 2012), and is available upon request.

SAVAGE is designed to construct individual haplotypes from the overlap graph of individual reads. We processed the totality of sequence data in a single patch in order to maximize the sensitivity of the contiguous reconstruction with parameter ‘--split 1’, as well as the parameter ‘--revcomp’ to handle reverse complement reads. The IPD exon 2 reference database that was used for MiSeq and DSR genotyping was also used to assess the quality of SAVAGE genotypes. The IPD sequences were mapped to contigs produced by SAVAGE using sensitive parameters in bbmap (minlength=100 vslow=t subfilter=0 indelfilter=0 lengthtag=t ignorefrequentkmers=t kfilter=100) designed to identify sequences that perfectly match SAVAGE contigs. A post-processing script refined these mappings, and only retained the mappings where the length of the mapped region was the same as the length of the IPD exon 2 sequence. These mappings indicated where the reference database sequence was fully and exactly contained within a SAVAGE contig.

## Results

### MHC reads are efficiently enriched using target-capture probes

We designed a custom target enrichment probeset that accounts for the extensive duplication of macaque MHC genes (Prall et al. 2017). A tripartite target capture system was used in this study. The first component is SeqCap EZ HGSC VCRome2.1, an optimized human clinical exome probeset. Since macaques are closely related to humans, SeqCap EZ HGSC VCRome2.1 can also be used in macaques, though some sequences that are most divergent between macaques and humans are not efficiently captured. The second component of the target capture system is an additional 22,884 rhesus macaque exon sequences that were not effectively captured with the SeqCap EZ HGSC VCRome2.1 reagent. The third component is a collection of probes designed to specifically enrich macaque MHC class I and class II sequences. These probes span the full length of HLA class I and class II genes including introns, 3’ UTRs, and approximately 1,000 bp of flanking 5’ sequence.

We obtained a median coverage of 100X for the target exon sequences across the genome with >20X coverage for 94.97% of bases that were targeted in this study. As illustrated in **Table 1**, an average of 70,549,789 Illumina sequence reads per sample were obtained for the 27 animals evaluated in this study. These reads were mapped against reference files containing representative genomic HLA exons 2 -3 for *HLA-A*, and *HLA-E* exons 2 - 4 for *HLA-DRA, -DRB1, -DRB3, -DRB4, -DRB5, -DQA1, -DQB1, -DPA1* and -*DPB1* sequences. We identified an average of 269,057 MHC class I and class II sequence reads per sample which corresponds to an average of 0.37% of the total sequence reads evaluated per sample (**Table 1**). In a previous study by Ericsen *et al*. (Ericsen et al. 2014), we found that MHC sequences only accounted for an average of 0.13% of the total Illumina sequence reads that were evaluated per animal when the standard HGSC VCRome2.1 panel was used alone for target capture (**Supplementary Table 1**). Thus, we achieved an almost three-fold increase of MHC genomic sequences after inclusion of the spike-in probes for target capture compared to use of the VCRome2.1 probeset alone.

**Table 1.**
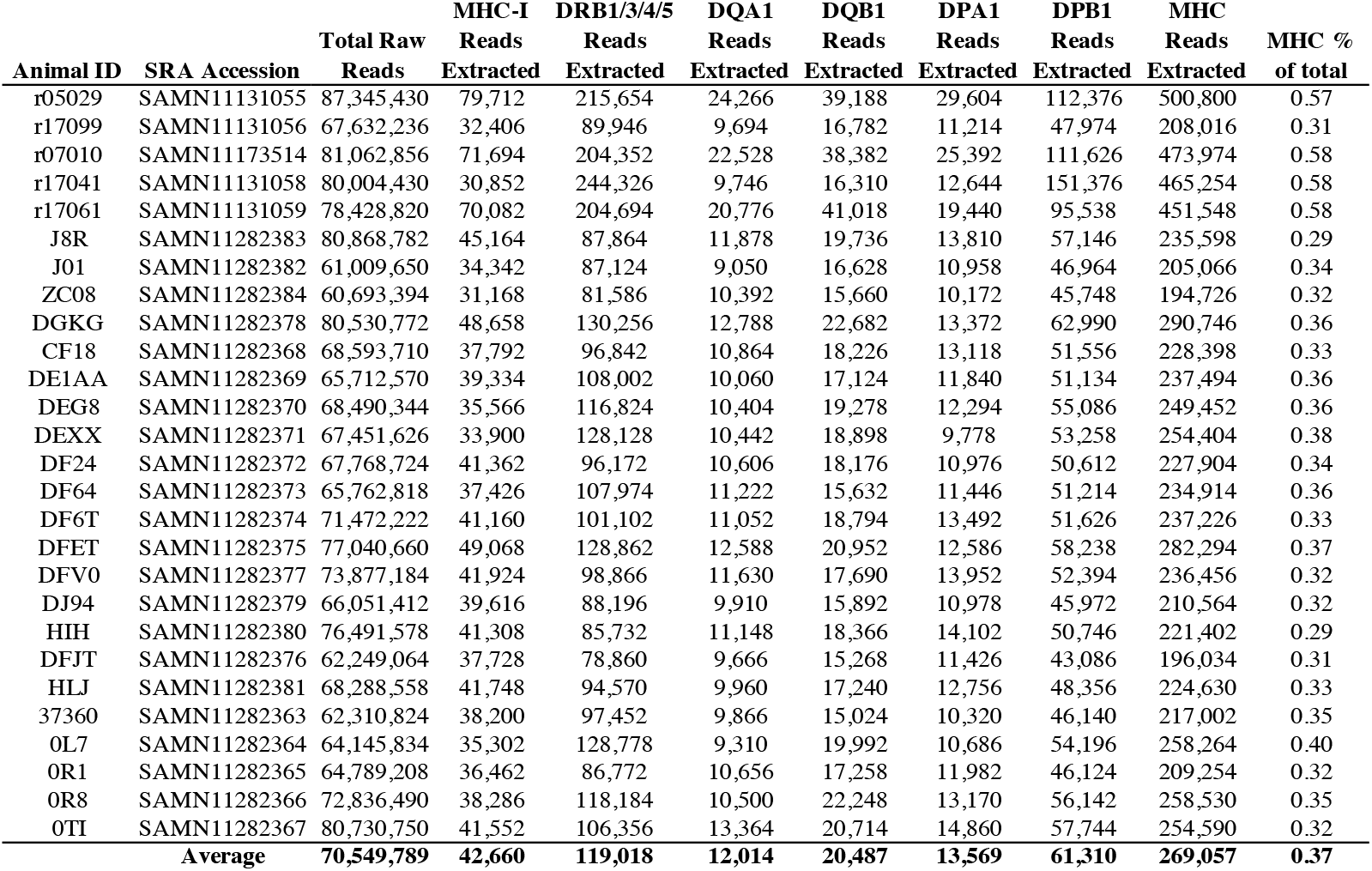
Fraction of total exome sequence reads corresponding to exons 2 - 3 of MHC class I and exons 2 - 4 of MHC class II genes.

### MHC genotypes defined from target-enriched genomic DNA are comparable to those obtained by amplicon deep sequencing

We hypothesized that MHC genotypes derived from target-enriched genomic sequence would be comparable in accuracy to MHC genotypes derived from conventional amplicon deep sequencing. MHC class I and class II PCR amplicons were generated from the same 27 animals and deep sequenced on an Illumina MiSeq. Representative MHC class I and class II genotypes supported by the amplicon data are shown in **Figure 1** and **Figure 2**, respectively. These figures illustrate genotypes from a pedigreed family of Indian rhesus macaques: a Sire, a Dam, and three Progeny that are paternal half-siblings. Comprehensive genotypes for all 27 Indian rhesus macaques evaluated in this study are shown in **Supplementary Figure 1**.

**Figure 1.**
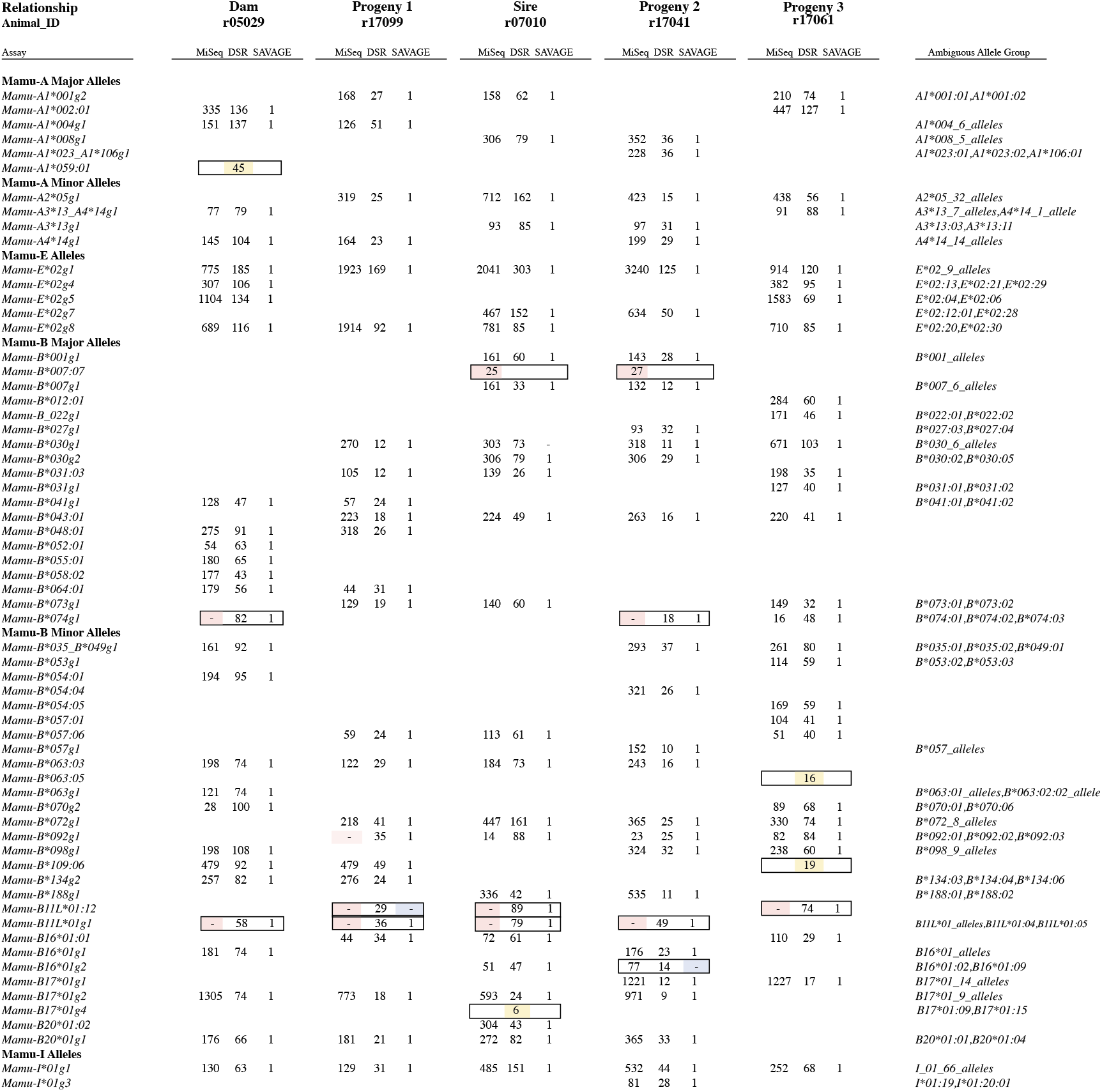
Comparison of MHC class I results from MiSeq PCR amplicon versus whole exome genotyping strategies for a representative breeding group of rhesus macaques. Results for each of the three methods are provided side-by-side in the columns for each macaque. Each row indicates the detection of a specific MHC class I allele or lineage group of closely related sequences that are ambiguous because they are identical over the IPD exon 2 database sequence. Values in the body of this figure indicate the number of sequence reads supporting each allele call for the MiSeq and DSR methods while alleles supported by a SAVAGE contig are reported with a “1”. Discrepancies between the MiSeq (pink), DSR (yellow) or SAVAGE (blue) methods are highlighted by filled cells with borders.

**Figure 2.**
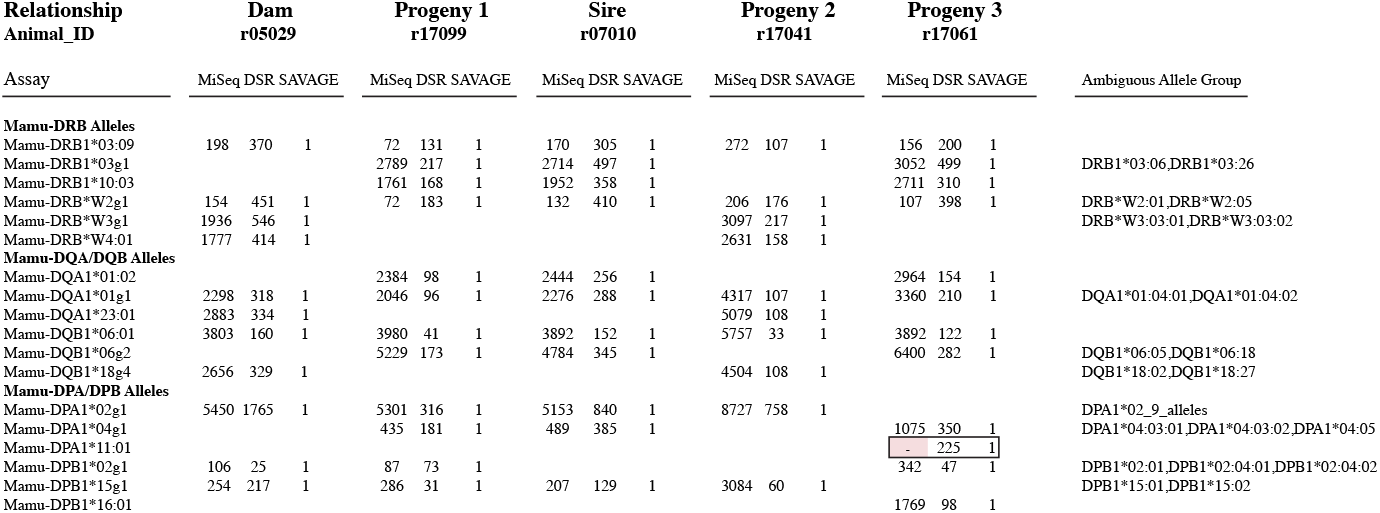
Comparison of MHC class II results from MiSeq PCR amplicon versus whole exome genotyping strategies for a representative breeding group of rhesus macaques. Results for each of the three methods are provided side-by-side in the columns for these five related macaques. Each row indicates detection of a specific MHC class II allele or lineage group of closely related sequences that are ambiguous because they are identical over the IPD exon 2 database sequence. Values in the body of this figure indicate the number of sequence reads supporting each allele call for the MiSeq and DSR methods while alleles supported by a SAVAGE contig are reported with a “1”. The *Mamu-DPA1*11:01* allele missed by the MiSeq assay due to multiple mismatches versus the amplification primers is highlighted in pink.

The superset of alleles supported by at least two of the methods described in this manuscript (MiSeq amplicon genotyping, DSR genotyping of individual exome sequencing reads, or SAVAGE genotyping of contigs derived from exome data), was considered to be the expected MHC genotypes for these animals. A complication for this assertion was certain genotypes were difficult for MiSeq analysis to report correctly. As we have shown previously (Karl et al. 2017), the number of reads supporting each genotyping call was highly variable, ranging from tens of reads to thousands of reads per allele. A small subset of allelic variants contained nucleotide substitutions in their sequences within the binding sites of the PCR oligonucleotides that interfere with efficient PCR amplification. This result is exemplified when members of the *Mamu-B11L*01* allele lineage were closely examined. Members of the *Mamu-B11L*01* allele lineage have two nucleotide substitutions relative to the 5’ oligonucleotide that was used to generate MHC class I amplicons, so these *Mamu-B11L*01* sequences were routinely absent in MiSeq genotypes (**Figure 1**). Likewise, the *Mamu-DPA1*11:01* allele has four nucleotide substitutions relative to the oligonucleotide pair used to generate *DPA1* amplicons for MiSeq genotyping (**Figure 2**).

These nucleotide substitutions do not fully account for all differences in the abundance of sequence reads for each allele. False negatives were also noted for certain allele lineages, such as *Mamu-B*074* and *Mamu-B*098*, which exhibited significantly diminished PCR efficiency despite being completely matched with the oligonucleotides used for amplification (**Figure 1**). False positive genotyping calls were also noted in the MiSeq assay that resulted from intermolecular recombination during the PCR process. This was exemplified by read support for the presence of *Mamu-B*007:07* in Sire r07010, and Progeny 2 r17041 (**Figure 1**). *Mamu-B*007:07* only differs from the *Mamu-B*007g1* allele group that was determined to be present in this pair of animals by a single nucleotide variant at the extreme 5’ end of the class I genotyping amplicon. Chimeric PCR products equivalent to the *Mamu-B*007:07* sequence were formed between the 3’ portion of the *Mamu-B*007g1* sequence and other allelic variants in these animals with this 5’ SNP. Taken together, these results illustrate that the MiSeq amplicon genotyping, while generally reflective of an animal’s MHC genotype, can yield both false positive and false negative results. When compared to the expected genotypes for this dataset, MiSeq amplicon genotyping has 90.4% MHC class I and 97.8% MHC class II concordance (**Table 2**).

**Table 2.**
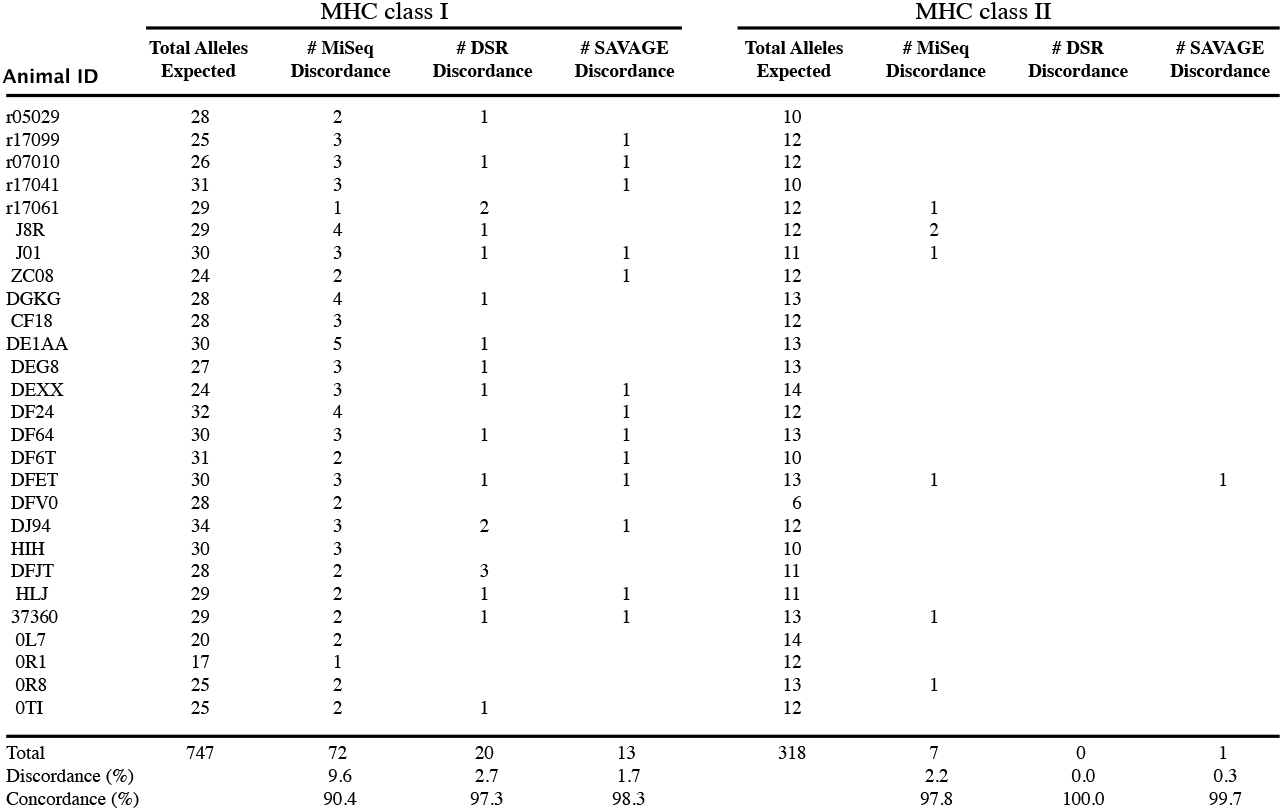
Summary of discordant results for each MHC genotyping method by animal.

Two separate strategies were used to derive MHC genotypes from target-capture data. The DSR analysis was a straightforward extension of the methodology used for deriving genotypes from MiSeq amplicons. MHC sequence reads were extracted from the total exome dataset. A simplified workflow that mapped those MHC reads to reference sequences, and then assigned the genotypes based on the presence of exome reads overlapping each position of the reference sequence, was problematic. This simplified workflow was extremely vulnerable to false positive genotypes, and the similarities among different MHC sequences often enabled those reads to map to multiple different alleles (Wiseman et al. 2013). Two or more reads each could partially match a portion of a sequence in the IPD exon 2 database, which could complement each other to provide support for an allele that was not biologically relevant. As discussed in the Methods, the DSR encompassed polymorphisms that discriminated among closely related alleles by requiring at least one mapped read to unambiguously map to the corresponding allele within the IPD exon 2 sequences. This strategy can identify specific polymorphisms of interest among the IPD exon 2 sequences to produce genotypes for MHC class I and MHC class II (**Figures 1, 2** and **Supplementary Figure 1**). For the 27 animals evaluated in this study, DSR was 97.3% and 100% concordant with expected MHC class I and class II genotypes, respectively (**Table 2**).

The DSR strategy did not consider sequences that were not among the IPD exon 2 sequences and this lack of consideration contributes to overcalled alleles. The apparent *Mamu-A1*059:01* allele in Dam r05029 was erroneously derived from reads that were from *Mamu-A1*004g1* and an unknown allele that was not among the sequences in the IPD exon 2 database. As a result of this unknown sequence not being among the known IPD exon 2 sequences, reads unambiguously mapped to *Mamu-A1*059:01*, which caused this allele to be overcalled. This type of overcalling will be mitigated with the discovery of additional allelic variants, and their inclusion in future iterations of the IPD database.

The second approach for determining genotypes performed *de novo* assembly on MHC class I and class II reads from each sample. The resulting assembled contigs were then mapped against IPD exon 2 reference sequences to define perfectly matching contigs. Because most assemblers were not tuned for the challenge of assembling large numbers of contigs that differ from one another by as little as 1 bp, we relied on an assembler, SAVAGE, originally designed to reconstruct viral sequencing haplotypes. As shown in **Figures 1, 2 and Supplementary Figure 1**, contigs produced by SAVAGE matched the expected genotypes. Unlike the DSR method, SAVAGE assembled contigs for downstream analyses. False positives and false negatives among closely-related variants were mitigated by the inclusion of the filtering steps described under Strategy 2. Across all 27 samples, the SAVAGE contigs are 98.3% and 99.7% accurate with respect to the expected MHC genotypes (**Table 2**).

While this analysis focused on genotyping using the same exon 2 reference sequences that are commonly utilized for MiSeq amplicon analyses, the SAVAGE contigs are frequently much longer than these reference sequences (**Figure 3**). These extended contigs frequently contain exons 2 through 4, plus the intervening introns, and could be used to provide higher resolution genotyping than is possible using exon 2 sequence alone. Moreover, contigs that contain complete sequences for exons 2-3 of MHC class I and exon 2 of MHC class II alleles meet the minimum criteria for obtaining formal allele nomenclature for non-human primates from the IPD-MHC (Robinson et al. 2013).

**Figure 3.**
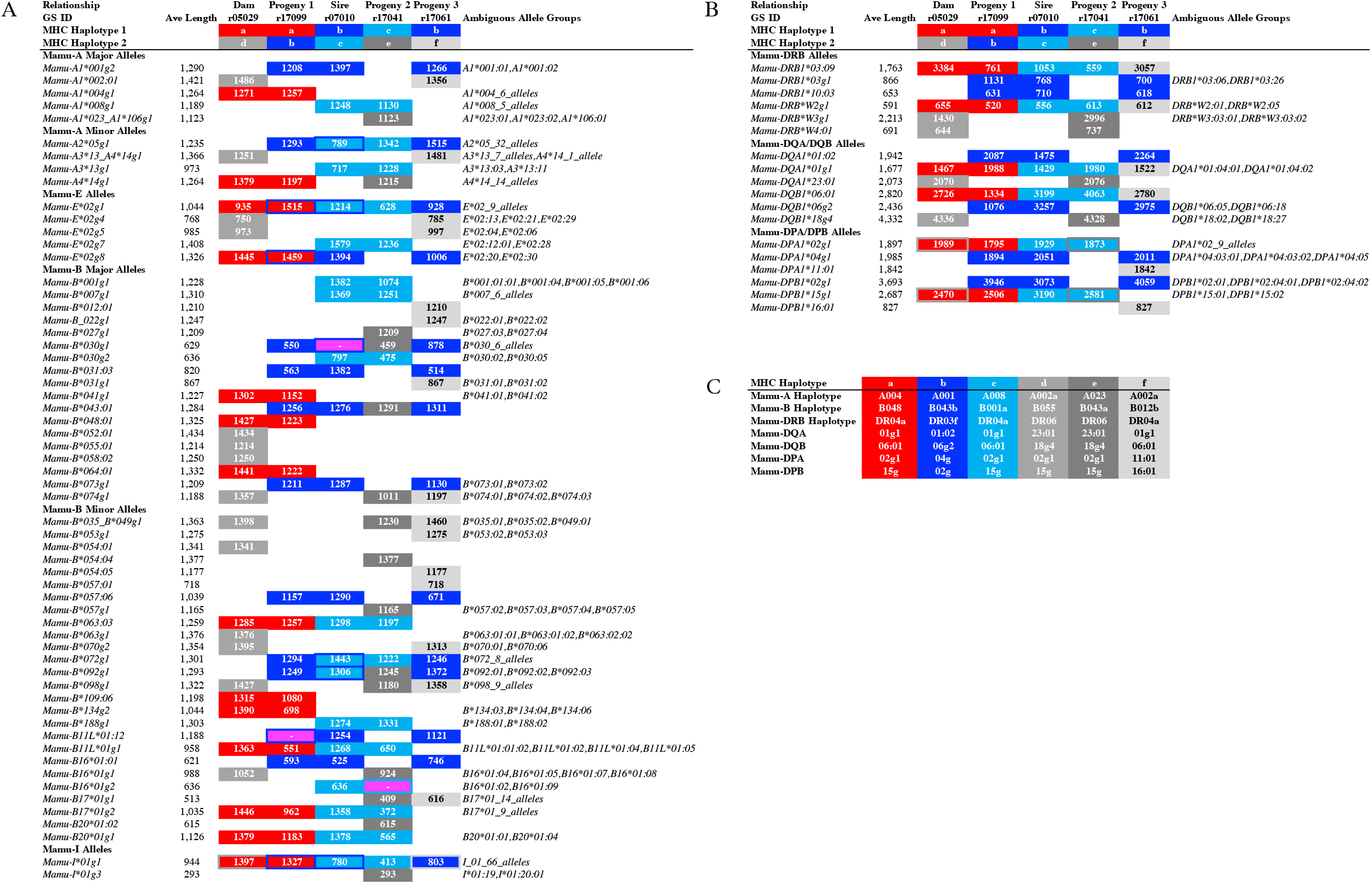
Lengths of MHC sequence contigs assembled by the SAVAGE method for a representative breeding group of rhesus macaques. Values in the body of this figure indicate contig lengths generated for these MHC sequences in each animal. (**A**) MHC class I genotyping results are illustrated for the five animals in this breeding group. Three false negative allele calls for SAVAGE versus the expected genotypes for these animals are highlighted in magenta, e.g., *Mamu-B*030g1* in Sire r07010. Sequences highlighted in red are associated with the maternal MHC Haplotype a that was inherited by Progeny 1 from its Dam. Sequences in Progeny 1 and Progeny 3 for their MHC Haplotype b that was inherited from Sire r07010 are highlighted in dark blue while the light blue sequences represent the alternate paternal Haplotype c that was inherited by Progeny 2. Three unique extended MHC Haplotypes in this breeding group are indicated with shades of grey (Haplotypes d - f). MHC allele groups that are shared by both parental haplotypes are indicated by colored borders around filled cells. (**B**) MHC class II genotyping results are illustrated for this same breeding group. (**C**) Abbreviated Mamu haplotype designations (Karl et al. 2013) are summarized for the six extended MHC haplotypes identified by segregation in this breeding group. For example, the twenty MHC sequences (red) that are associated with extended Haplotype a can be summarized by the following string of abbreviated Mamu haplotype designations: Mamu-A004/B048/DR04a/DQA01g1/DQB06:01/DPA02g1/DPB15g.

## Discussion

Here we describe MES genotyping of Indian rhesus macaques. In our estimation, this method will supersede MiSeq PCR amplicon genotyping as the most widely used macaque MHC genotyping assay in the future. This methodology has a number of compelling advantages. Most importantly, exome sequencing dramatically improves the overall quantity and quality of genomic information obtained from each sample. The same datasets used for MHC analyses may be used to evaluate protein-coding genetic variation throughout the genome. The loss of start and stop codons exome-wide can be obtained from the same datasets by modifying the workflows *in silico*, as opposed to requiring the sequencing of multiple new sets of target genes (Yang et al. 2013). Exome-wide datasets offer the promise of retrospectively identifying candidate DNA sequence variants that may be responsible for unexpected experimental outcomes in studies with macaques and other nonhuman primates. The availability of exome-wide sequences in conjunction with MHC genotypes may also increase the rigor of prospective macaque experiments by enabling more sophisticated balancing of experimental groups, and exclusion of animals whose genetics are likely to strongly bias experimental results (Loffredo et al. 2007; Reynolds et al. 2011; Haus et al. 2014).

These same exome datasets also offer the potential to improve the quality of MHC genotyping. Full-length long-read MHC transcript sequencing offers exquisite resolution, but this technology can be labor intensive and difficult to scale (Karl et al. 2017; Semler et al. 2018). The MiSeq exon 2 amplicon approach is limited in its allelic resolution, but it is the current standard deep sequencing approach for high-throughput MHC genotyping in macaques. In this report, we compare two novel strategies for MHC genotyping from MES datasets to this standard MiSeq amplicon genotyping method. The results that we obtained with all three approaches show strong concordance with expected MHC genotypes for the Indian rhesus macaques that were evaluated. As illustrated in **Figure 3a**, MHC class I genomic contigs with an average length of approximately 1.1 kb can be assembled from sequence reads that were initially extracted from the whole exome datasets.

Following the initial enrichment step, MHC class II genomic contigs averaging approximately 1.9 kb in length could be assembled from whole exome sequence reads that mapped to *HLA-DRB1, -DRB4, -DRB5, -DQA1, - DQB1, -DPA1* and -*DPB1* reference sequences, which contained exons 2 - 4, and the pairs of introns (**Figure 3b**). The addition of HLA spike-in probes containing both exon and intron sequences for the target capture step, and the introduction of 151 bp paired end reads for the Illumina NovaSeq platform greatly facilitated our ability to assemble these extended MHC genomic contigs. Additional technological advances, including even longer sequence reads and more efficient assembly algorithms, will undoubtedly increase genomic contig lengths as well as MHC allelic resolution in future studies.

Both MHC genotyping approaches described here depend upon mapping exome sequence reads against a reference database of MHC class I and class II allele sequences. Currently, the IPD database essentially is restricted to coding regions for macaque MHC sequences, and many IPD entries are only partial transcript sequences that lack complete coding regions. In addition, it is very challenging to correctly phase short Illumina sequence read that map to different exons of a specific allelic variant and are separated by intronic sequences that span hundreds to thousands of base pairs in genomic DNA. Our results with the SAVAGE workflow (**Figure 3**) demonstrate that exome sequence reads that have been enriched with the enhanced MHC probe design described here can be assembled into genomic contigs that span multiple exons and introns. These contigs, therefore, could also be used to improve MHC reference databases, though this requires a major effort with more animals, and is beyond the scope of this manuscript.

The current cost of genotyping is relatively high compared to amplicon deep sequencing, largely due to two expenses. First, the amount of sequence data needed for a single sample can be high. Compared to conventional MHC genotyping, where 192 macaques’ data can be collected on a single instrument run, exome data acquisition is much more expensive. On an Illumina Novaseq instrument, which has a much higher run cost than the MiSeq, only 70 exome samples can be sequenced simultaneously per lane of a S4 flow cell. However, this cost of sequencing is rapidly decreasing, and as of early 2019, commercial providers have advertised sequencing for $9 USD per Gb of whole genome sequence data^1^. Second, a major expense in MHC genotyping is the production, validation, and use of target-capture arrays, as well as the development of *in silico* data analysis workflows. The approaches described here, in particular MHC genotyping from SAVAGE contigs produced by *de novo* assembly of exome reads, is flexible and should be adaptable as sequencing approaches evolve and improve.

A major goal for future studies will be to attempt to extend these contigs to encompass full length genomic MHC sequences using SAVAGE or other assembly software tools. The relatively compact genomic structure and consistent length of MHC class I genes increase the attainability of this goal. Establishment of comprehensive macaque MHC allele databases of extended genomic sequences will greatly facilitate mapping of exome sequence reads since they will be contiguous with the reference sequences instead of being interrupted by intervening sequences between each exon that are not included in current non-human primate IPD-MHC databases (Maccari et al. 2017).

These results demonstrate that MHC genotypes can be obtained by analyzing genomic DNA selectively enriched for MHC and protein-coding gene sequences. This represents an important advance for characterizing MHC genetics in macaques, and this suggests that analyses of whole exome and whole genome data will become the predominant method for studying macaque genetics in the coming decade.

## Acknowledgements

We gratefully acknowledge Michele Di Mascio and his group at the National Institute of Allergy and Infectious Diseases of the National Institutes of Health for providing rhesus macaque samples used in this study. We also gratefully thank Brian Bushnell for assistance with the BBTools software, and the WNPRC for providing samples from five related macaques.

This research was supported by contracts HHSN272201600007C from the National Institute of Allergy and Infectious Diseases of the National Institutes of Health. This work was also supported in part by the Office of Research Infrastructure Programs/OD (P51OD011106) awarded to the Wisconsin National Primate Research Center at the University of Wisconsin-Madison. This research was also supported in part by grant R24-OD011173 from the National Institutes of Health. This research was conducted in part at a facility constructed with support from Research Facilities Improvement Program grants RR15459-01 and RR020141-01.

## Supplementary Materials

**Supplementary Table 1.** Fraction of total exome sequence reads corresponding to MHC class I and class II genes after target capture with the VCRome2.1 probe design alone. This exome sequence dataset was described previously by Ericsen and coworkers (Ericsen et al. 2014).

| GS ID | SRA Accession | Total Raw Reads | MHC-I Reads Extracted | DRB1/3/4/5 Reads Extracted | DQA1 Reads Extracted | DQB1 Reads Extracted | DPA1 Reads Extracted | DPB1 Reads Extracted | MHC Reads Extracted | MHC % of total |
| --- | --- | --- | --- | --- | --- | --- | --- | --- | --- | --- |
| cy0642 | **SAMN11281734** | 79,795,404 | 4,848 | 26,822 | 2,568 | 2,274 | 900 | 81,340 | 118,752 | 0.15 |
| cy0643 | **SAMN11281735** | 86,265,844 | 4,408 | 26,930 | 2,542 | 2,320 | 936 | 74,370 | 111,506 | 0.13 |
| cy0646 | **SAMN11281736** | 109,909,444 | 4,724 | 25,476 | 2,382 | 2,196 | 910 | 72,206 | 107,894 | 0.1 |
| cy0648 | **SAMN11281737** | 106,315,304 | 5,558 | 34,442 | 2,966 | 2,714 | 1,092 | 97,254 | 144,026 | 0.14 |
| cy0649 | **SAMN11281738** | 78,575,038 | 6,022 | 33,742 | 3,176 | 2,970 | 1,134 | 90,556 | 137,600 | 0.18 |
| cy0651 | **SAMN11281739** | 102,890,862 | 6,686 | 33,194 | 3,388 | 3,094 | 1,294 | 94,532 | 142,188 | 0.14 |
| cy0652 | **SAMN11281740** | 89,074,544 | 4,652 | 25,314 | 2,660 | 2,478 | 976 | 69,976 | 106,056 | 0.12 |
| cy0654 | **SAMN11281741** | 99,169,798 | 5,294 | 30,328 | 2,924 | 2,756 | 1,042 | 79,664 | 122,008 | 0.12 |
| **Average** |  | **93,999,530** | **5,274** | **29,531** | **2,826** | **2,600** | **1,036** | **82,487** | **123,754** | **0.13** |

**Supplementary Figure 1.** Comparison of MHC class I results from MiSeq PCR amplicon versus whole exome genotyping assays with the DSR and SAVAGE strategies for all 27 animals. Results for each of the three method are provided side-by-side in the columns for each macaque. Each row indicates the detection of a specific MHC class I allele or lineage group of closely related sequences that are ambiguous because they are identical over the IPD exon 2 database sequence. Values in the body of this figure indicate the number of sequence reads supporting each allele call for the MiSeq and DSR methods while alleles supported by a SAVAGE contig are reported with a “1”. Discrepancies between the MiSeq (pink), DSR (yellow) or SAVAGE (blue) methods are highlighted by filled cells with borders.

## Footnotes

1 Comment via Twitter 03/18/2019 @albertvilella https://twitter.com/AlbertVilella/status/1107524501645000705

